# Natural Genetic Variation in *Drosophila melanogaster* Reveals Genes Associated with *Coxiella burnetii* Infection

**DOI:** 10.1101/2020.05.21.109371

**Authors:** Rosa M. Guzman, Zachary P. Howard, Ziying Liu, Ryan D. Oliveira, Alisha T. Massa, Anders Omsland, Stephen N. White, Alan G. Goodman

## Abstract

The gram-negative bacterium *Coxiella burnetii* is the causative agent of Query (Q) fever in humans and coxiellosis in livestock. Association between host genetic background and *Coxiella burnetii* pathogenesis has been demonstrated both in humans and animals; however, specific genes associated with severity of infection remain unknown. We employed the *Drosophila* Genetics Reference Panel to perform a genome-wide association study and identify host genetic variants that affect *Coxiella burnetii* infection outcome. The analysis resulted in 64 genome-wide suggestive (P < 10^−5^) single nucleotide polymorphisms or gene variants in 25 unique genes. We examined the role of each gene in *Coxiella burnetii* infection using flies carrying a null mutation or RNAi knockdown of each gene and monitoring survival. Of the 25 candidate genes, 15 validated using at least one method. For many, this is the first report establishing involvement of these genes or their homologs with *Coxiella burnetii* susceptibility in any system. Among the validated genes, *FER* and *tara* play roles in the JAK-STAT, JNK, and decapentaplegic/TGF-β signaling pathways that are associated with the innate immune response to *Coxiella burnetii* infection. Two other two validated genes, *CG42673* and *DIP-ɛ*, play roles in bacterial infection and synaptic signaling but no previous association with *Coxiella burnetii* pathogenesis. Furthermore, since the mammalian ortholog of *CG13404* (*PLGRKT*) is an important regulator of macrophage function, *CG13404* could play a role in *Coxiella burnetii* susceptibility through hemocyte regulation. These insights provide a foundation for further investigation of genetics of *Coxiella burnetii* susceptibility across a wide variety of hosts.

## INTRODUCTION

*Coxiella burnetii* is the causative agent of Q fever, a zoonotic disease that poses a serious threat to both human and animal health (Maurin and Raoult, 1999). Based on morbidity, low infectious dose and the environmental stability of the organism, *C. burnetii* is classified as a Category B priority pathogen by the United States NIH and CDC (Madariaga et al., 2003). It is well known that the epidemiology of Q fever is associated with the presence of infected animals; sheep, goats, cattle, and humans primarily become infected by inhalation of contaminated aerosols (Marrie et al., 1996; McQuiston et al., 2002; Schimmer et al., 2008). Therefore, reducing bacterial load in the livestock is critical to preventing Q fever outbreaks. *C. burnetii* is endemic worldwide and sporadic outbreaks have recently been reported in the United States (Anderson et al., 2013; Dahlgren et al., 2015; Karakousis et al., 2006; Kersh et al., 2013; Sondgeroth et al., 2013). A recent large outbreak of Q Fever that originated in a goat farm in the Netherlands provides a warning of the risks associated with *C. burnetii* infection where 307 million Euros were spent in public health management efforts and agricultural interventions (Roest et al., 2011a, 2011b; Schimmer et al., 2008; van Asseldonk et al., 2013). To date, no commercial Q fever vaccine is available for humans or animals in the United States, and antibiotic therapy is the only option for treating the infection in humans. Culling infected or at-risk animals has been a strategy to contain emerging outbreaks (Roest et al., 2013, 2011b, 2012). Additionally, the lack of animal models with genetic malleability and the strict requirements for BSL3 animal facilities for work with Select Agent phase I virulent strains of *C. burnetii* make it difficult to study host-pathogen interactions *in vivo*.

The host genetic background has been shown to influence the development of *C. burnetiid* infection in both humans and other animals (De Lange et al., 2014; Delaby et al., 2012; Ghigo et al., 2002; Leone et al., 2004; Meghari et al., 2008; Raoult et al., 2005). Experimental studies in human and mouse cells have correlated defective monocyte/macrophage activation and migration with ineffective granuloma formation and overexpression of IL-10 observed in patients with chronic Q fever (Bewley, 2013; Delaby et al., 2012; Ka et al., 2014; Meghari et al., 2008; Mehraj et al., 2013). Two recent studies performed genotyping in human population and revealed that genetic variation in innate immune genes, such as pattern recognition receptors and *IFNG*, are associated with susceptibility to Q fever (Ammerdorffer et al., 2016; Wielders et al., 2015). Despite this importance, additional unidentified factors and specific genetic variants associated with susceptibility to the infection remain largely unknown. In addition, it remains to be elucidated how host genetic factors affect bacterial load and shedding in susceptible reservoir hosts.

Previous studies have profiled mammalian host responses to *C. burnetii* infection and show that the bacteria down-regulate the host innate immune response during acute infection and that the resolution of Q fever is associated with the re-establishment of type I interferon signaling (Faugaret et al., 2014; Ghigo et al., 2002; Gorvel et al., 2014). However, the mechanisms by which *C. burnetii* targets the host innate immune pathways and the host genes that contribute to the block in innate immune activation remain largely unknown. Directed studies in humans have revealed that single-nucleotide polymorphisms (SNPs) in innate immune receptors and signaling genes such as *TLR1, STAT1, IFNG*, and *MyD88* are associated with acute or chronic Q fever (Schoffelen et al., 2015; Wielders et al., 2015). Since these studies used a targeted approach to examine SNPs in only a set of candidate genes, we aim to undertake a global, genome-wide analysis to identify gene variants associated with *C. burnetii* infection using *Drosophila melanogaster* as the host model.

We recently demonstrated that adult *D. melanogaster* flies are susceptible to infection with the BSL2 NMII clone 4 strain of *C. burnetii* and that this strain is able to replicate in this host (Bastos et al., 2017). Additionally, *D. melanogaster* lacking functional copies of the gram-negative bacteria sensing immune deficiency pathway (Imd) signaling genes, *PGRP-LC* and *Relish*, displayed increased susceptibility to *C. burnetii* infection. Indeed, the *D. melanogaster* model is suitable for studying host-pathogen interactions during *C. burnetii* infection. Importantly, the recently developed *Drosophila* genetics reference panel (DGRP), a fully sequenced, inbred panel of fly lines derived from a natural population, provides an efficient platform for genotype-to-phenotype associations via a genome-wide association study (Huang et al., 2014a; Mackay et al., 2012a). The DGRP has already been used to reveal genes associated with resistance/tolerance to other bacterial pathogens (Bou Sleiman et al., 2015; Howick and Lazzaro, 2017; Wang et al., 2017).

In this study, we identified genetic variants in *D. melanogaster* that were associated with susceptibility or tolerance to *C. burnetii* infection. Specifically, we obtained a list of 64 SNPs in 25 unique genes from the different GWA performed. Our analyses revealed genes with sex-specific effects on susceptibility and tolerance that have functions associated with actin binding, transcriptional response, and regulation of G-proteins. Multiple genes within the decapentaplegic (DPP) pathway, homologous to the TGF-β pathway in mammals (Gelbart, 1989), were associated with susceptibility to infection. Rho GEFs and TGF-β have been associated with the development of the *Coxiella*-containing vacuole (CCV) and pathogenesis in humans, respectively (Aguilera et al., 2009; Benoit et al., 2008c, 2008a; Pennings et al., 2015; Salinas et al., 2015; Weber et al., 2016). Importantly, all the candidate genes identified have mammalian orthologs or highly conserved functions that will allow for extrapolation of the findings here to mammalian systems.

Of the 25 candidate genes identified, 15 genes significantly affected host survival during *C. burnetii* infection in *D. melanogaster* null mutants or RNAi knockdown flies. While other DGRP studies tend to use either null mutants or RNAi knockdown flies for validation studies, we used both gene disruption methods to test how the genes affect diseases severity. We also examined the effect of candidate SNPs using regulatory element analysis (modENCODE) and found some within transcription factor binding hot spots, putative enhancers, novel splicing, branch point variation, and codon usage variation that could explain how the variants may affect host gene expression ultimately altering the ability to fight infection. We show how the DGRP can be utilized to identify host genetic variants associated with sex-specific susceptibility or tolerance to *C. burnetii* infection, and have broad cross-species applications.

## MATERIAL AND METHODS

### *Drosophila melanogaster* and *C. burnetii* stocks

Fly stocks were obtained from the Bloomington *Drosophila* Stock Center, the Vienna *Drosophila* Resource Center, Exelixis at Harvard Medical School and the Kyoto Stock Center. Fly stocks were maintained at room temperature in standard meal agar fly food at 25°C and 65°C humidity. All fly strains used are listed in **Table S1**. *C. burnetii* Nine Mile phase II (NMII) clone 4 RSA439 was propagated in Acidified Citrate Cysteine Medium 2 as previously described (Omsland et al., 2009). *C. burnetii* stocks were quantified by measuring bacterial genome equivalent (GE) using quantitative real-time PCR (qPCR) as previously described (Coleman et al., 2004).

### Fly infections and hazard ratio phenotype determination

Groups of 40 male and female adult flies (two to seven days old) from each DGRP line used in this study (**Table S2 and S3**) were injected with phosphate buffered-saline (PBS) or 10^5^ genome equivalents (GE) of *C. burnetii* diluted in PBS to establish infection. For injections, flies were anesthetized with CO_2_ and injected with 23 nL of bacteria or PBS using a pulled glass capillary and an automatic nanoliter injector (Drummond Scientific, Broomall, PA). Individual flies were injected at the ventrolateral surface of the fly thorax and placed into new vials. After injection, mortality was monitored daily for 30 days with the flies maintained at 25°C and 68% humidity. Survival curves were analyzed by the log-rank (Mantel-Cox) test using GraphPad Prism (GraphPad Software, Inc.) to determine a hazard ratio for males and females for each DGRP line. Any line having less than three percent mortality in the mock-infected group was not included in downstream analyses.

### Genome-wide association analyses using hazard ratios

Phenotype to genotype association was performed by submitting log_10_ transformed hazard ratios to the dgrp2 webtool (http://dgrp2.gnets.ncsu.edu/), which adjusts the phenotype for the effects of *Wolbachia* infection and major inversions (Huang et al., 2014b; Mackay et al., 2012b). Three separate analyses were run using male, female, and combined hazard ratios for the DGRP lines used in this study (**Table S2-S4**). R was used to create QQ plots from log-transformed hazard ratios and obtain R^2^ values and genomic inflation values (λ). For male and female analyses, 193 male and 195 female log-transformed hazard ratios were submitted (**Table S2 and Table S3**, respectively). SNPs and small indel variants with a P-value (mixed effects model) below 10^−5^ were considered genome-wide suggestive candidate SNPs and further analyzed. For the combined analysis, both male and female hazard ratios were submitted for 191 DGRP lines. Candidate SNPs with P-values (mixed effects model) less than 10^−5^ for the average trait or the difference (female – male) trait were selected for further study. The DGRP genome assembly (BDGP R5/dm3) was used to identify variants in candidate genes. Human orthologs of candidate genes were identified by using the DRSC Integrative Ortholog Prediction Tool (DIOPT) and the ortholog with the highest weighted score was reported (Hu et al., 2011a). Predicted functions for each candidate gene were gathered using Flybase (FB2019_02). Regulataory annotation summaries for each SNP and indel were compiled using Flybase (FB2019_02) and modENCODE utilizing variant coordinates converted to the BDGP R6/dm6 reference assembly. Regulatory annotations were expounded by reviewing publicly available data within modENCODE tracks, all noncoding features including transcription factor binding sites, histone ChIP-seq data, chromatin domain segmentation, and small RNA-seq tracks.

### Validation of candidate genes

Two replicates of forty adult flies from each null mutant and RNAi knockdown for each candidate gene were injected with PBS or *C. burnetii*, as stated previously, to empirically determine the effect of knockout or knockdown of the candidate gene on severity to infection. All experiments were conducted twice independently, and the results were combined to determine hazard ratios and generate mortality graphs. RNAi knockdown was performed using straight-winged progeny from crosses between the CyO-balanced *Act5C-GAL4* driver line and the corresponding dsRNA-containing RNAi lines (**Table S1**). Sibling progeny flies carrying the CyO balancer were used as control flies. Genetic background strains for each null mutant strain were used as control flies. After injection, adult flies were maintained at 25°C and 68% humidity for 30 days and mortality was monitored daily. Hazard ratios were determined from the survival curves of two combined mortality experiments and analyzed as stated previously. In addition, we employed a strict threshold of P < 0.01 to determine significant change from control genotype as summarized in **Table S5-6**.

### Splicing, branch point variation, and codon usage analysis

The Ensembl project (http://uswest.ensembl.org/index.html) and The Human Splicing Finder (http://www.umd.be/HSF/) were used to determine splicing and branch point variation from curated sequences to determine codon usage fraction based on frequency of amino acids per thousand.

### Data availability

Strains and stocks are available upon request. Genomic sequence for the DGRP is available at http://dgrp.gnets.ncsu.edu/. Supplemental material is available at FigShare. The authors affirm that all data necessary for confirming the conclusions of the article are present within the article, figures, and tables.

## RESULTS

### Susceptibility to *C. burnetii* infection is dependent on host genetic background

Previously, we determined that flies deficient in the IMD signaling pathway genes, *PGRP-LC* and *Relish*, exhibited increased susceptibility to *C. burnetii* infection (30). We also determined that the gene *eiger* contributed to decreased tolerance to *C. burnetii* infection in flies, and *eiger* mutant flies were less susceptible to *C. burnetii* infection (Bastos et al., 2017). Therefore, we hypothesized that susceptibility to *C. burnetii* infection in *Drosophila* was associated with host genetic background, and that the broad base genetic variation in the DGRP would help identify other candidate genes that effect susceptibility to *C. burnetii* infection via GWA analysis. To determine the susceptibility of each DGRP line to *C. burnetii* infection, adult males and females of each line were mock-infected or infected with *C. burnetii*. Hazard ratios were calculated using the survival curves for male and female flies and subsequently used as input for the GWA analysis (**Figure 1**). In total, 193 and 195 hazard ratios were calculated for males and females, respectively. The survival curves revealed an approximately log normal distribution of hazard ratios ranging from −0.719 to 1.643 for male flies (0.191 to 44.01, non-log-transformed) and −0.714 to 1.200 for female flies (0.1932 to 15.85, non-log-transformed) (**Table S2 and S3, Figure S1A**), which indicates that genetic polymorphisms in the DGRP lines affect susceptibility to *C. burnetii* infection. Interestingly, male flies were more susceptible than female flies overall to *C. burnetii* infection, with a mean hazard ratio of 1.90 for male flies and 1.56 for female flies (P = 0.0015) (**Figure S1A**). Notably, we observed three distinct susceptibility phenotypes for both male and female flies. Most DGRP lines were susceptible or resistant to *C. burnetii* infection, characterized by increased mortality in the *C. burnetii* group or similar mortality in the *C. burnetii* and mock-infected group, respectively. However, some lines exhibited decreased mortality compared to the mock-infected group. Mutualistic traits maintained through the evolution of *C. burnetii* may act synergistically with specific genotypes in the DGRP lines to yield a symbiotic phenotype.

**Figure 1.**
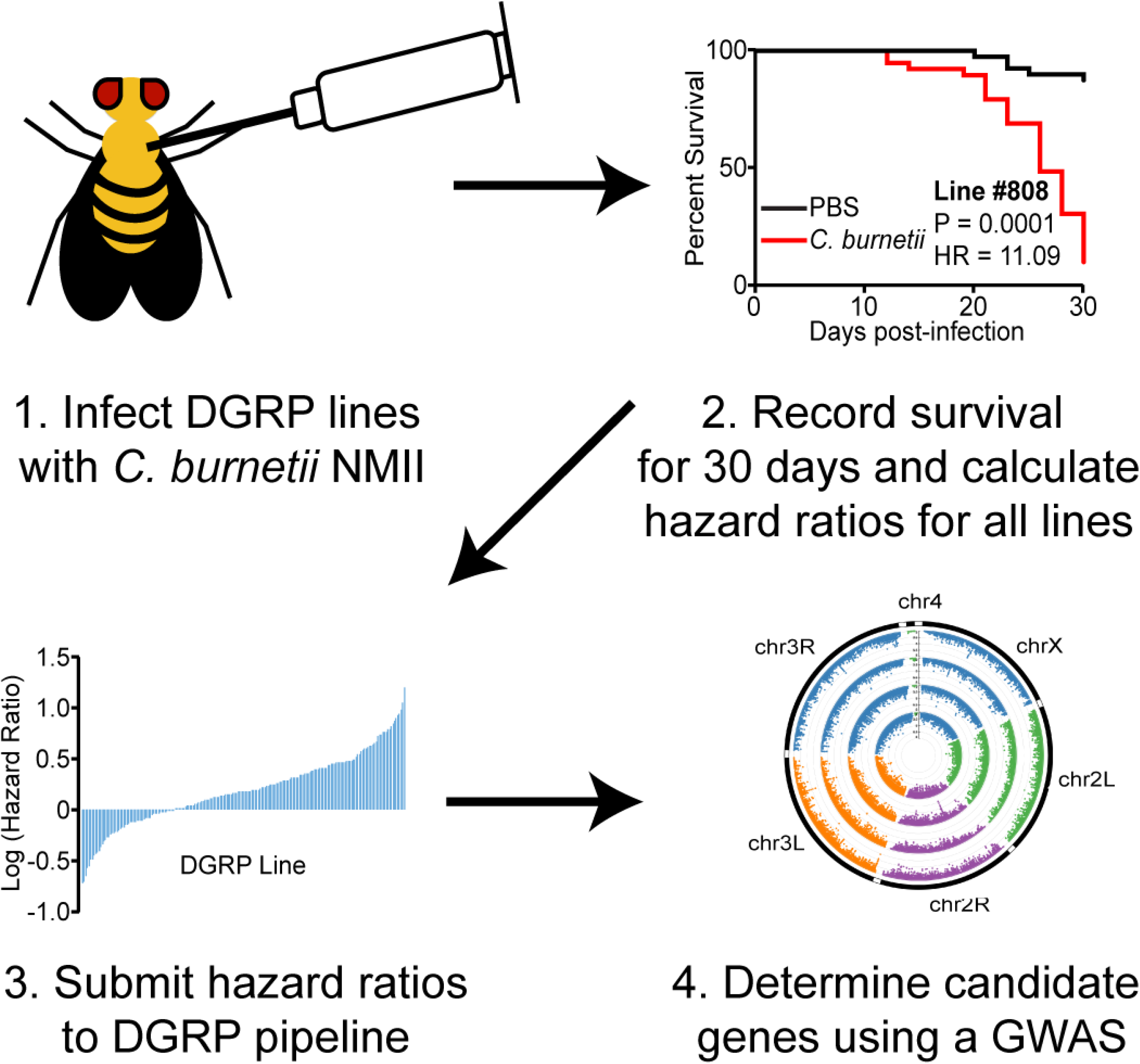
Experimental design schematic. Groups of 40 males and females per DGRP line were injected with PBS or *C. burnetii* at 10^5^ bacteria/fly and host survival monitored for 30 days to obtain hazard ratios. The hazard ratios of all DGRP lines were log-transformed and used as input for a GWAS.

**Figure 2.**
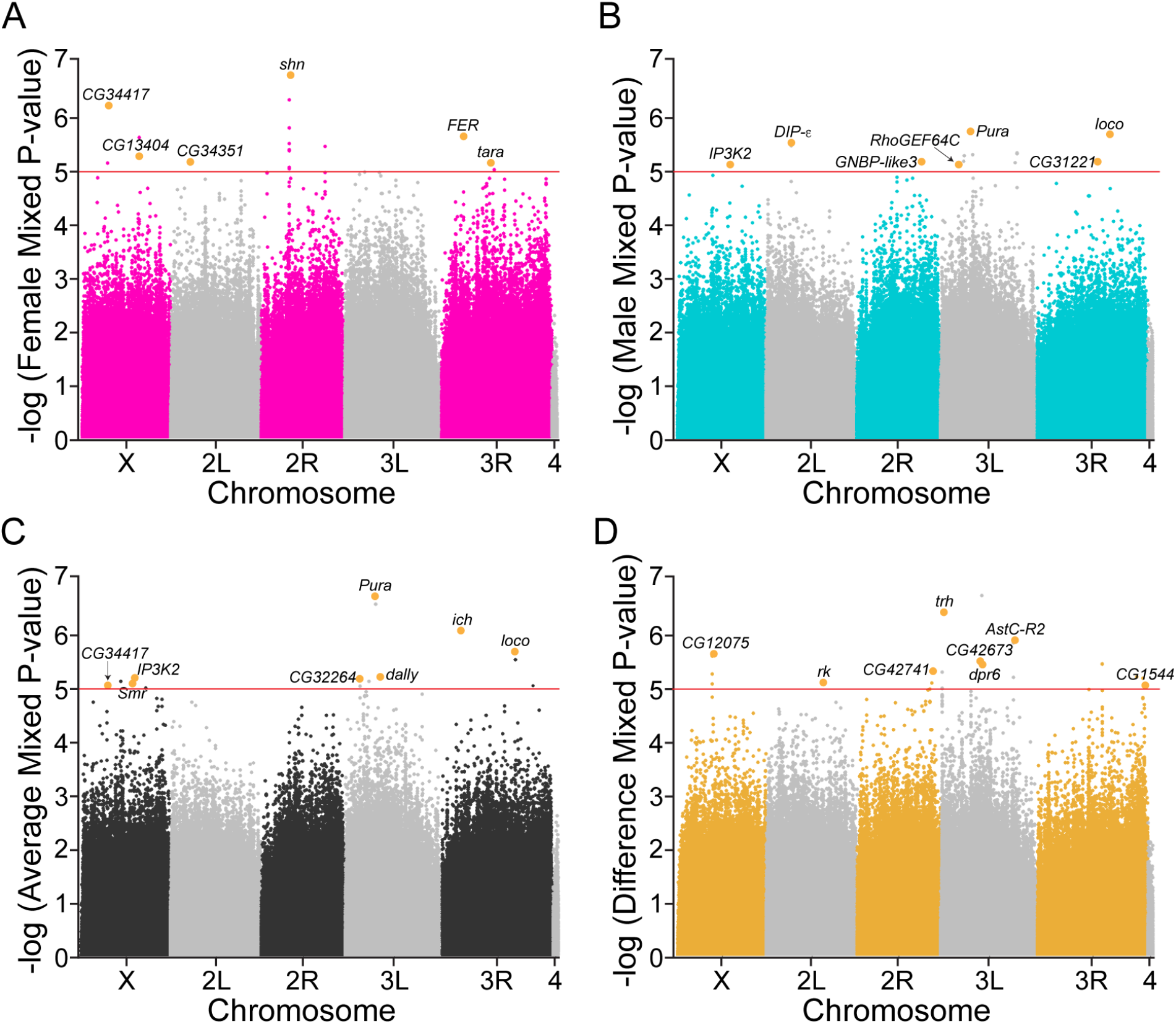
Genome-wide association analyses with hazard ratios reveal candidate genes. Manhattan plots for A) male, B) female, C) average, and D) difference GWA analyses using mixed effect P-values for all four traits from dgrp2 webtool. Highlighted SNPs with P-values below 10^−5^ are labeled with associated candidate genes.

### GWA analyses of DGRP hazard ratios reveals candidate SNPs

The DGRP facilitates rapid genome-wide association analyses using a quantitative phenotype via submission of a data set to the online webtool (Mackay et al., 2012a). To determine polymorphisms in the DGRP population that affect susceptibility to *C. burnetii*, the hazard ratios were submitted for analysis. However, due to genome-wide association analyses relying on parametric tests, the hazard ratios were log transformed to yield an approximately normal distribution (Shapiro-Wilke test, P > 0.1) prior to submission for GWA analysis (**Figure S1A**). Additionally, it was determined that hazard ratios were significantly positively correlated between male and female flies (P = 4.99 × 10^−7^), but with an r^2^ value of 0.121, which indicates a weak correlation and potential sex-dependent genotypes (**Figure S1B**). Thus, hazard ratios were submitted as separate files for male and female analyses, and a single, combined file in order to identify polymorphisms that may be sex-dependent and to increase power for polymorphisms that are sex-independent. The sex-independent analysis which we termed average analysis, results in top hit SNPs that affect both sexes while the sex-dependent analysis which we termed difference analysis, results in top hits that affect one sex but not the other. A total of 193, 195, and 191 hazard ratios were submitted for males, females, and average and difference, respectively (**Table S2-S4**).

A total of 1,893,791 polymorphisms were tested in the male analysis, and 1,897,049 polymorphisms were tested in the female analysis. The combined analysis tested a total of 1,889,141 polymorphisms. In total, the GWA analyses yielded a total of 64 associated polymorphisms, of which 59 were unique polymorphisms, below the genome-wide suggestive P-value threshold of 10^−5^, as expected for studies employing the DGRP (**Table S5**). In addition, quantile-quantile (Q-Q) plots revealed no significant inflation due to dataset distribution, and lambda values ranged from 0.993 (females) to 1.002 (difference) (**Figure S2A-D**). Lastly, P-values derived from these analyses appear to be reduced overall based on the lines from the Q-Q plots and lambda values below 1 (**Figure S2A-D**).

Of the 64 polymorphisms identified from the GWA, 14 SNPs are intergenic (21.9%), three of which are within 200 base pairs upstream of nearest gene; 39 are within introns (60.9%); eight are within exons (12.5%); one is within the 5’ UTR (1.6%); and two are in antisense-coding RNA within exon/introns (3.1%) (**Table S5**). Of the eight SNPs within exons, six are silent and two are missense mutations. From the 64 top SNPs, we identified 25 unique candidate genes with available stocks for gene disruption and used the DGRP genome assembly (BDGP R5/dm3) to gather predictive functions and regulatory annotations for each gene using Flybase (FB2019_02) and modENCODE (**Table 1**). For each candidate gene we also report the human ortholog with the highest weighted score from the DRSC Integrative Ortholog Prediction Tool (DIOPT).

**Table 1.** Candidate genes associated with top SNPs from GWA analyses.

| Candidate Gene | Top SNP (BDGP R5/dm3) | P-value | Analysis | Regulatory Annotations | Putative Gene Function | Human Ortholog* |
| --- | --- | --- | --- | --- | --- | --- |
| *CG34351* | 2L_4702261_SNP | 7.5x10<sup>-6</sup> | Female | Poorly annotated<br>Euchromatin transcriptionally silent (or intergenic) | Regulation of G-proteins | RGS7BP |
| *DIP-ε* | 2L_6394872_SNP | 2.9x10<sup>-6</sup> | Male | Euchromatin transcriptionally silent or dynamic | Interaction with Dprs | OPCML |
| *rk* | 2L_13999491_SNP | 8.4x10<sup>-6</sup> | Difference | Transcriptionally silent | GPCR; buriscon receptor; melanization | LGR5 |
| *shn* | 2R_7099616_SNP | 1.9x10<sup>-7</sup> | Female | TFBS hot spot | Zinc finger C2H2 transcription factor | HIVEP2 |
| *GNBP-like3* | 2R_16414194_SNP | 6.1x10<sup>-6</sup> | Male | Euchromatin transcriptionally silent or dynamic | Beta 1,3-glucan recognition/binding | CRYBG1 |
| *CG42741* | 2R_18904195_SNP | 5.3x10<sup>-6</sup> | Difference | Transcriptionally silent | Zinc finger C2H2 transcription factor | KLF3 |
| *trh* | 3L_376337_SNP | 4.4x10<sup>-7</sup> | Difference | TFBS | bHLH-PAS transcription factor | NPAS1 |
| *CG32264* | 3L_3750617_SNP | 7.5x10<sup>-6</sup> | Average | Transcriptionally silent | Actin binding | PHACTR2 |
| *RhoGEF64C* | 3L_4738164_SNP | 7.4x10<sup>-6</sup> | Male | Euchromatin transcriptionally silent or dynamic | Rho guanyl-nucleotide exchange factor | ARHGEF3 |
| *Pura* | 3L_7623383_SNP | 2.2x10<sup>-7</sup> | Average | lncRNA | Rho guanyl-nucleotide exchange factor | PLEKHG4 |
| *dally* | 3L_8851042_SNP | 6.7x10<sup>-6</sup> | Average | Putative enhancer but not hot spot | Co-receptor for growth factors/morphogens | GPC5 |
| *CG42673* | 3L_9540740_SNP | 3.5x10<sup>-6</sup> | Difference | TFBS | Nitric-oxide synthase binding | NOS1AP |
| *dpr6* | 3L_10044744_SNP | 3.8x10<sup>-6</sup> | Difference | Transcriptionally silent | Interaction with DIPs | CADM1 |
| *AstC-R2* | 3L_18481371_SNP | 1.4x10<sup>-6</sup> | Difference | Between two TFBS | Allatostatin receptor | SSTR2 |
| *ich* | 3R_4787301_SNP | 9.3x10<sup>-7</sup> | Average | TFBS hot spot | Zinc finger C2H2 transcription factor | PRDM15 |
| *FER* | 3R_5218712_INS | 2.7x10<sup>-6</sup> | Female | Active enhancer | Protein tyrosine kinase activity | FER |
| *tara* | 3R_12079260_SNP | 7.7x10<sup>-6</sup> | Female | Active enhancer, TFBS hot spot | Transcriptional co-regulator | SERTAD1 |
| *CG31221* | 3R_15278653_SNP | 6.4x10<sup>-6</sup> | Male | Near TFBS | LDL receptor | LRP1B |
| *loco* | 3R_18456211_SNP | 2.3x10<sup>-6</sup> | Average | Antisense RNA, enhancer, TFBS | Regulation of G-proteins | RGS12 |
| *CG1544* | 3R_27026419_SNP | 9.4x10<sup>-6</sup> | Difference | TFBS | Oxoglutarate dehydrogenase | DHTKD1 |
| *CG34417* | X_6434578_SNP | 9.9x10<sup>-6</sup> | Average | TFBS hot spot, putative enhancer/promoter | Actin binding | SMTN |
| *CG12075* | X_8751630_SNP | 2.7x10<sup>-6</sup> | Difference | Silent chromatin state | Lipid signaling | - |
| *Smr* | X_12610055_SNP | 8.9x10<sup>-6</sup> | Average | TFBS hot spot | Chromatin binding; transcriptional regulation | NCOR1 |
| *IP3K2* | X_13210675_SNP | 7.0x10<sup>-6</sup> | Average | Putative enhancer site | Calcium regulation; IP3 signaling | NET1 |
| *CG13404* | X_14160126_SNP | 6.1x10<sup>-6</sup> | Female | Active enhancer, TFBS hot spot | Plasminogen receptor (KT) | PLGKRT |

\*human orthologs from DIOPT. Ortholog with highest weighted score reported.

We analyzed the chromatin state of all 25 candidate gene variants using Flybase and modENCODE and found that 12 are in transcription factor binding sites (TFBS) (48%); nine are within regions predicted to be transcriptionally silent (36%); one is within a long noncoding RNA (4%); and three are in enhancers only (12%) (**Table 1**). The SNP within the candidate gene *loco* lies within an antisense RNA that is also an enhancer and TFBS. Candidate gene functions were derived from available information on Flybase.

### Validation of candidate genes

We next tested the 25 candidate genes from the different GWA (**Table 1**) by infecting and monitoring survival during *C. burnetii* infection for 30 days in flies carrying a null mutation in the candidate gene or knocked down for the candidate gene by RNAi. We defined validation of candidate genes as any line that has significantly different mortality than the mock-infected and genetic control (**Tables S6 and S7**). Specifically, we employed a strict threshold of P < 0.01 to determine significant change from control genotype. Of the 25 candidate genes, six validated in null mutants only (24%), five in RNAi knockdown only (20%), four in both null mutants and RNAi (16%), and 10 did not validate with either method (40%) (**Figure 3A**). Mortality of *w^1118^* males and females (**Figure 3B-B’**) during *C. burnetii* infection were used as the genetic control for several null mutants, including *RhoGEF64C^MB04730^* (**Figure 3C**), *tara^1^* (**Figure 3D**), and *CG13404^f07827b^* (**Figure 3E**). We selected these candidate genes to represent how validation was determined based on P-value and survival trend for validating genes from different categories, i.e. null-only, RNAi-only, or both. *w^1118^* females (**Figure 3B**) are not susceptible to *C. burnetii* infection (P = 0.033) but *w^1118^* males (**Figure 3B’**) are highly susceptible (P < 0.0001) which corroborates our previous work (Bastos et al., 2017). The candidate gene *RhoGEF64C^MB04730^* was selected from male-only GWA (**Figure 3C-C’**) and we observed that survival in null mutants (**Figure 3C**) was overall tolerant (P = 0.0014) compared to *w^1118^* males (**Figure 3B’**). In contrast, there was no significant change in survival between control and RNAi genotypes (**Figure 3C’**) (control, P = 0.0374; RNAi, P = 0.0130). Thus, *RhoGEF64C^MB04730^* males validated only in null mutants. The candidate gene *tara* was selected from the female-only GWA and we observed that in null mutants (**Figure 3D**) and RNAi knockdown flies (**Figure 3D’**), the absence of the gene resulted in increased mortality compared to control genotypes. Specifically, *tara*^1^ females are susceptible to infection (P < 0.0001) compared to *w^1118^* females (**Figure 3B**) and *tara* RNAi females (**Figure 3D’**) are also susceptible (P = 0.0025) compared to control (P = 0.0123). Thus, *tara* validated for females in both null mutants and RNAi knockdown flies. The candidate gene *CG13404 ^f07827b^* was selected from female-only GWA and we observed that null mutants (**Figure 3E**) are not susceptible to infection (P = 0.2737) like *w^1118^* females (**Figure 3B**). In contrast, *CG13404* RNAi females (**Figure 3E’**) are susceptible to infection (P < 0.0001) while control genotype females are not (P=0.3914). Thus, *CG13404* validated only in RNAi knockdown flies.

**Figure 3.**
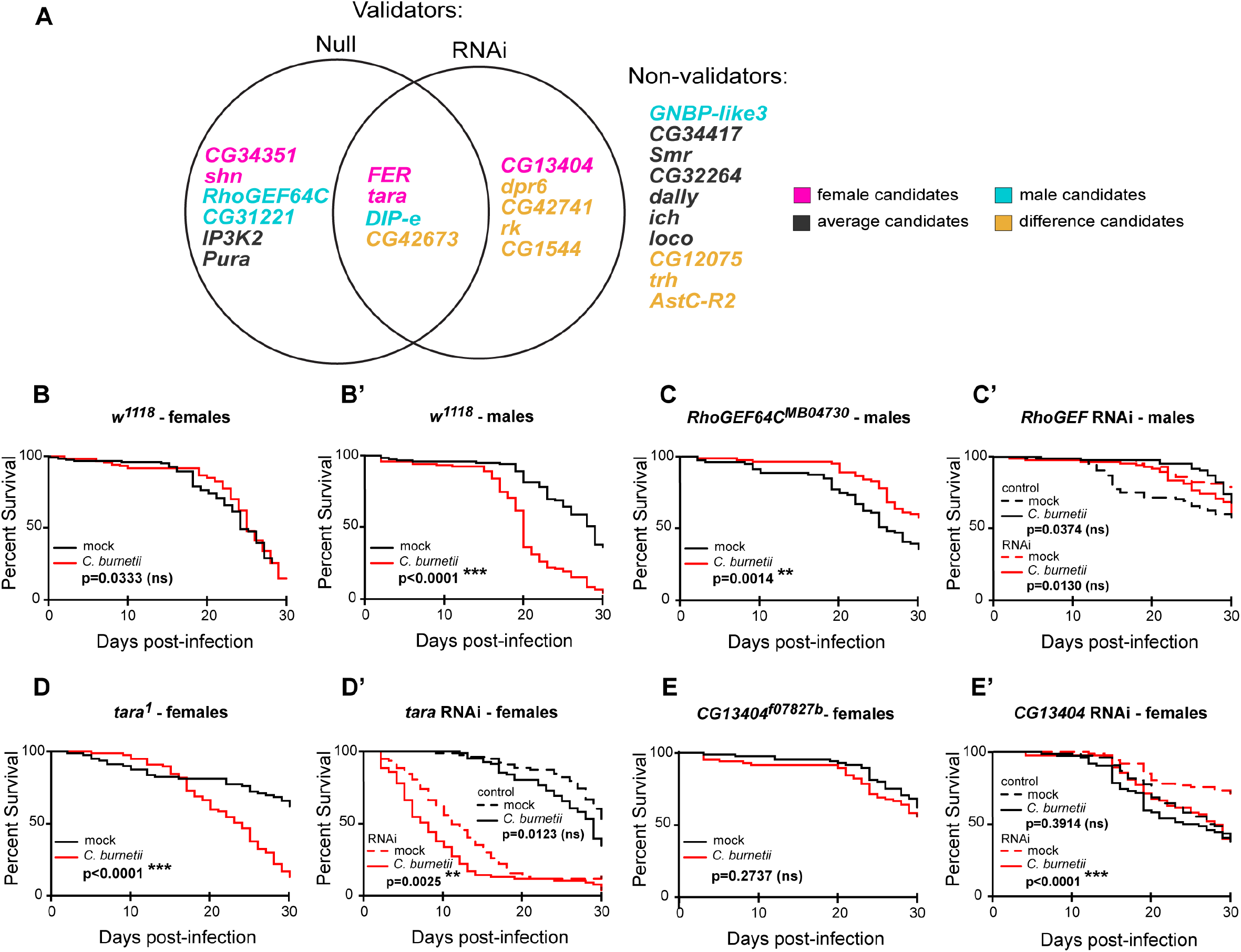
Fifteen GWAS candidate genes are validated using survival rate as a metric. (A) Venn diagram summarizes the genes that validate in null mutant flies, RNAi knockdown flies, or both. A gene is validated if the P value for its survival curve (mock versus infected) changes between the control and experimental line. P value threshold were not significant (n.s.), P < 0.01, and p < 0.0001. Colors indicate the type of GWAS analysis from which the gene came. (B-B’) Survival curves of control *w^1118^* (B) females and (B’) males following mock or *C. burnetii* infection. (C-E) Survival curves of *RhoGEF64C^KG02832^* (C) or control and *RhoGEF64C* RNAi males (C’), *tara^1^* (D) or control and *tara* RNAi females (D’), or *CG13404^f07827a^* (E) or *CG13404* RNAi females (E’) following mock or *C. burnetii* infection. Each survival curve represents two independent experiments of at least 40 flies that were combined for a final survival curve, Statistical significance (Log-rank test) from the mock-infected group is indicated.

### ENCODE analysis of validated genes

Splicing and branching of precursor mRNA and abundance of tRNA codons are known to affect gene expression (Jeacock et al., 2018; Komar, 2016; Královičová et al., 2004; Sauna and Kimchi-Sarfaty, 2011; Singh and Cooper, 2012; Wang and Burge, 2008; Will and Luhrmann, 2011; Zhou et al., 2016). Therefore, we used data available from the ENCODE project to determine regulatory annotations for the SNPs in genes that validated in host survival experiments. **Table 2** summarizes the splicing and branch point analysis in terms of percent variation from wildtype and codon usage as a fraction of frequency of amino acid (SNP) per thousand over frequency of amino acid (wildtype) per thousand. Several SNPs varied at the predicted mRNA splicing sites, branch points, or codon usage compared to wildtype sequences such as the SNPs affecting the validated genes *CG34351*, *DIP-ɛ*, *Pura*, *tara*, *FER*, and *IP3K2*. Changes in condon usage fraction for *shn* and *CG13404* may affect these gene variants albeit to a lesser extent. Together, we use these results to hypothesize why gene variants may or not validate using null mutants and RNAi knockdown.

**Table 2.** Splice, branch point, and codon usage analysis of validating genes.

| Candidate gene | Top SNP (BDGP R5/dm3) | Splice Variation From Wildtype | Branch Point Variation From Wildtype | Codon change | Codon Usage Fraction (wildtype/SNP)* |
| --- | --- | --- | --- | --- | --- |
| *CG34351* | 2L_4702261_SNP | No difference | -34.14% | - | - |
| *DIP-ε* | 2L_6394872_SNP | N/A | N/A | Ctg/Ttg (silent) | 3.72 |
| *shn* | 2R_7099616_SNP | N/A | N/A | ttA/ttG (silent) | 1.1 |
| *Pura* | 3L_7623460_SNP | 7.74% | No difference | - | - |
| *tara* | 3R_12079260_SNP | 64.82% | N/A | - | - |
| *FER* | 3R_5218712_INS | -92.69% | No difference | - | - |
| *IP3K2* | X_13210675_SNP | -17.01% | No difference | - | - |
| *CG13404* | X_14160126_SNP | N/A | N/A | ctC/ctT (silent) | 0.86 |

\*Frequency of amino acid per thousand

## DISCUSSION

Host genetics are known to lead to differences in human Q fever severity, and previous studies using human samples were focused on specific genes such as *IFNG, STAT1, and TLR10* (Ammerdorffer et al., 2016; Wielders et al., 2015). In this study, we performed an unbiased GWAS to uncover all possible genes that may regulate *C. burnetii* infection. Our study builds on previous literature by identifying host variants that not only affect *C. burnetii* infection outcome but can also be translated to future human studies due to conserved sequences. In addition to host variants, bioinformatic analysis of regulatory elements also reveals further possible explanations for host survival differences between genotypes. Altogether, the validated genes in this study reveal both novel connections and conserved cross-species connections like hemocytes/macrophage regulation and sex-specific susceptibility differences.

To perform the GWAS we infected *D. melanogaster* lines from the DGRP library, which have known genetic variation, with *C. burnetii* and measured host survival. We then used the hazard ratios from the survival curves as input for a genome-wide association to identify host variants affecting host survival. In the genetic screen alone, over 31,000 individual fruit flies were infected yet additional power of the DGRP screen lies on the previously genetically annotated 4,565,215 naturally occurring molecular variants, including 3,976,011 high quality single/multiple nucleotide polymorphisms (SNPs/MNPs), 125,788 polymorphic microsatellites, 169,053 polymorphic insertions and 293,363 polymorphic deletions (relative to genome reference) (Mackay and Huang, 2018). Our screen revealed 64 genome-wide suggested polymorphisms, 0.0014% of the 4,565,215 total variants.

We then narrowed the 64 SNPs to 25 unique candidate genes to test host survival during *C. burnetii* infection based on fly stocks available for gene disruption using null mutants and RNAi knockdown. We validated candidate genes using two rules; 1) That mutants (null or RNAi-knockdown) would have a statistically significant difference in survival compared to mock at a threshold of P < 0.0001, and 2) That the survival trend would differ from control genotype. Of the 25 candidate genes tested, 15 validated using either null mutants or RNAi knockdown. We did not expect that validating genes would phenotypically behave the same between null mutants and RNAi knockdown because the loss versus decreased levels of a gene may influence pathways in differently (Boettcher and McManus, 2015; Zimmer et al., 2019). Several *D. melanogaster* GWA studies use flies in which gene expression has been silenced using RNAi knockdown flies to validate candidate genes because controls for knockdown are obtained within sibling progeny of a cross (Howick and Lazzaro, 2017; Palu et al., 2019, 2020). In contrast, null mutants must be matched to a genetic control line (Chow et al., 2013; Swarup et al., 2013). Our results showed that while four candidate genes validated using both methods, 11 other candidate genes validated using one method of gene disruption (**Figure 3A**). These results suggest the need for rigor in that multiple methods should be employed when applicable for testing host survival or validating gene candidate (Ayroles et al., 2015).

Two major connections between the validated fly genes in this study and mammalian systems is the role of immune cell regulation and sex-specific differences. The gene *CG13404*, which was identified in the female-only GWA, can explain both of these themes. Host mortality in *CG13404* RNAi-knockdown flies indicated they were significantly more susceptible to infection compared to control genotype. The human ortholog of *CG13404* is the plasminogen receptor (*PLG-R_KT_)*, which is important for macrophage polarization and efferocytosis, two key components of inflammation regulation (Vago et al., 2019). The absence of *PLG-R_KT_* causes defective plasminogen binding and inflammatory macrophage migration in both male and female mice pups, but only female *Plg-R_KT_ ^−/−^* pups die two days after birth (Miles et al., 2017). We hypothesize that these sex-specific differences are conserved across species given that female flies knocked down for *CG13404* were more susceptible to *C. burnetii* infection.

In *D. melanogaster* immunity, hemocytes are the professional phagocytic cells given their ability to recognize, engulf, and destroy dying cells during development and pathogens during larval and adult stages (Hoffmann, 2003; Regan et al., 2013; Yano et al., 2008). Hemocytes also mediate the secretion of antimicrobial peptides (AMPs) in response to pathogen infection through the Toll, JAK/STAT, and Immune deficiency (Imd) pathways (Hoffmann, 2003; Lemaitre and Hoffmann, 2007). Recent studies in our lab have shown that hemocytes support *C. burnetii* replication and induce Imd-specific AMPs (Bastos et al., 2017; Hiroyasu et al., 2018). Finally, hemocytes play important roles in melanization, encapsulation, and coagulation/clotting (Vlisidou and Wood, 2015). For example, embryonic hematopoesis involves migration of progenitor blood cells and cytoskeleton rearrangement that requires integrins, Rho family GTPases, microtubule proteins, and actin/actin-binding proteins (Comber et al., 2013; Evans et al., 2010; Huelsmann, 2006; Paladi, 2004; Stramer et al., 2010; Zanet et al., 2009). While it remains undetermined how these individual processes occur during *C. burnetii* infection, our genetic screen identified genes that could be involved in haemocyte regulatory processes such as the *Rho guanyl-exchange factor* (*RhoGEF*) gene and two putative actin-binding proteins in the uncharacterized genes *CG34417* and *CG32264*. Future mechanistic studies are underway to determine the role of validated genes in the context of *C. burnetii* infection and will likely reveal more cross-species immune response conservation.

Additional connections to mammalian pathways can be extrapolated from the genes that validated in both null mutants and RNAi knockdowns, i.e. *DIP-ɛ, FER, tara*, and *CG42673*. The human orthologs of *DIP-ɛ, FER, tara*, and *CG42673*, are *OPCML, FER, SERTAD1*, and *NOS1AP*, respectively, based on highest weighted gene from the DRSC Integrative Ortholog Predictive Tool (DIOPT) score (Hu et al., 2011b). In flies, *tara* encodes a transcriptional co-regulator that interacts with chromatin remodeling complexes, cell cycle proteins, and the JNK signaling pathway and plays a role in ataxin-1-induced degeneration (Afonso et al., 2015; Branco et al., 2008; Calgaro et al., 2002; Fernandez-Funez et al., 2000). The ortholog *SERTAD1* is also a transcriptional co-regulator and has been linked to molecular neural abnormalities similar to *tara* (Biswas et al., 2010; Savitz et al., 2013). Interestingly, a recent study showed that induction of *SERTAD1* is IFN-independently expressed during Nipah virus infection (Glennon et al., 2015). It is known that IFN induction is tissue-dependent during *C. burnetii* infection (Hedges et al., 2016); therefore, it is plausible that *tara* is targeted during *C. burnetii* infection. We observed that the loss of *tara* lead to significantly decreased host survival during *C. burnetii* infection and future mechanistic studies on this gene may reveal novel host-pathogen interactions.

In flies, *DIP-ɛ* belongs to the immunoglobulin superfamily (IgSF) of *defective proboscis extension response* (Dpr) and *Dpr-interacting proteins* (DIP), which form a complex network of cell surface receptors in synaptic specificity. We observed that both null mutants and RNAi knockdown flies exhibited increased mortality to *C. burnetii* infection compared to their respective controls. We also noted that codon frequency fraction compared to wildtype is 0.27, which suggests that the SNP change results from decreased abundance of tRNA codon availability. How *DIP-ɛ* affects host survival at the cellular level remains unknown but for the SNP (2L_6394872), it may be due to decreased codon availability altering proper gene expression that is ultimately important to the fly to fight infection. We hypothesize that *DIP-ɛ* may play a novel role during infection.

*FER*, which leads to activation of DPP and is a TGF-β homolog, has recently been shown to improve survival of *Klebsiella pneumoniae* through *STAT3* when overexpressed (Dolgachev et al., 2018; Li et al., 2019; Murray, 2006). It is known that *C. burnetii* induces expression of *STAT3* and *IL-10* during murine infection (Millar et al., 2015; Murray, 2006; Textoris et al., 2010). One of these studies showed that male mice have increased gene expression of *STAT3* and *IL-10* during infection which may account for the higher susceptibility of Q fever observed in men (Textoris et al., 2010). Our study corroborates previously observed sex-specific differences in gene expression following *C. burnetii* infection given that *FER* was a top hit in the female-only GWA. We hypothesize that the absence of *FER* in females disrupts the immune response required to control infection and leads to significantly increased host mortality.

The last of our validating genes in both null mutants and RNAi knockdown is *CG42673*, an uncharacterized gene that was a top SNP (3L_9540740) hit in the difference GWA. Another DGRP GWAS found that loss-of-function of *CG42673* in blood cells significantly impaired the cellular immune response to *Staphylococcus aureus* (Nazario-Toole, 2016). Interestingly, this study also found that *dpr10* significantly affected *S. aureus* phagosome maturation while our own top hit, *dpr6* (3L_10044744_SNP) validated in RNAi knockdown. *CG42673* may function as an enhancer like its human ortholog *NOS1AP* and is perhaps a target of *C. burnetii* pathogenesis. *NOS1AP* is known to bind neural nitric oxide synthase in the brain as well as proteins involved in the spliceosome and small nucleolar ribonucleic complexes according to mass-spectrometry protein-interaction studies (Grossmann et al., 2015; Hein et al., 2015).

Another theme of this study is the role of regulatory elements among validated genes. The eukaryotic gene expression process depends on proper splicing of precursor mRNA into mature mRNA which requires spliceosome recognition at specific RNA sequences at the exon-intron boundaries (Singh and Cooper, 2012; Wang and Burge, 2008). Together with the 3’ and 5’ splice sites, the branch site helps bind small nuclear ribonucleic proteins for efficient exon recognition and branch point variants can result in exon skipping, aberrant splicing and altered production of transcripts that ultimately cause disease (Královičová et al., 2004; Will and Luhrmann, 2011). Similarly, the relative abundance of codons encoding each amino acid affects mRNA translation/stability and transcription ultimately determining gene expression levels (Jeacock et al., 2018; Komar, 2016; Sauna and Kimchi-Sarfaty, 2011; Zhou et al., 2016). We determined regulatory annotations for SNPs in genes that affected host survival using data available in ENCODE to reason how the mutations identified by the GWA can be affecting host survival, such as by affecting transcript abundance. We found that for several SNPs in validating genes differ in splicing, branch point variation, and codon abundance compared to wildtype sequences.

For example, the candidate gene *CG34351* was selected from female-only GWA and host mortality to *C. burnetii* infection was significantly different from genetic control in null mutants only (**Figure 3A, Table S5**). We found that the SNP (2L_4702261) within this gene differs −34.14% from wildtype branch point splicing. This large negative percentage difference suggests that the branch point site is broken. The SNP in validated gene *DIP-ɛ* (2L_6394872) showed a codon usage fraction of 3.72, higher than would be expected for an equal usage ratio of differing codons. We also found that the SNP (3L_7623460) within *Pura* differs 7.74% from wildtype splice variation, which indicates a new splice site creation with no destruction. The SNP (3R_12079260) within *tara* differs 64.82% from wildtype splicing which also indicates a new splice site creation with no destruction. For this *tara* SNP we interpret that the new acceptor site could end the intron at alternate site and could contribute to mortality of null mutants and RNAi knockdown flies. The insertion (3R_5218712) within *FER* differs −92.69% from wildtype splicing and has no variation in branch point splicing from wildtype. The −92.69% change in splicing for this *FER* SNP indicates the site is broken but the destruction is offset by a 3 base pair insertion. At the host level, *FER* null mutants and RNAi knockdowns differed from control genotypes, which may be partially explained by altered splicing. The SNP (X_13210675) within *IP3K2* differs −17.01% from wildtype splicing and has no variation in branch point splicing from wildtype. The −17.01% splicing difference in this *IP3K2* SNP indicates the splice site is broken because there is destruction with no creation.

At the regulatory annotation level, a silent codon change (ttA/ttG) was found in the SNP for *shn* (2R_7099616) (**Table 2**). However, in total, 10 unique SNPs were located within *shn* from the female-only GWA (**Table S4**), this gene validated in the null mutant but not by RNAi knockdown. Specifically, the *shn^1^* null mutant exhibited lower mortality during *C. burnetii* infection than the *y^1^ w^67c23^* genetic control, which we defined as a tolerant phenotype (**Table S5**). The human ortholog of *shn* is *HIVEP2* (HIV enhancer binding protein 2) which also encodes a transcription factor that binds to NF-κB of different genes and contains a zinc finger C2H2 transcription factor domain (Allen and Wu, 2005). Although no published studies have linked *HIVEP2* or *shn* to *C. burnetii* pathobiology, *shn* is part of the decapentaplegic (DPP) pathway, analogous to the human TGF-β pathway (36). Additionally, it has been shown that *C. burnetii* induces expression of TGF-β1 in during atypical macrophage maturation (Benoit et al., 2008b). There were no significant changes in splicing, branch point variation, or codon usage in the *shn* SNP compared to wildtype therefore we suspect this gene functions with others during *C. burnetii* infection or within a pathway not tested here. Overall, our results show that employing select bioinformatic tools allows us to browse integrative-level annotations on candidate and validated host variants in this study. Taken together with the host survival of variants during *C. burnetii* infection, several hypothesis-driven questions can be posed about the immune function of the variants.

In addition to connections between validated fly genes and their mammalian orthologs, there were interesting observations between the DGRP predictive effect of top hits and validating genes. According to the GWAS data, the *RhoGEF64C* SNP (3L_4738164) causes increased susceptibility (effect = −0.1709, **Table S5**). However, we observed that survival of *RhoGEF64C^MB04730^* null mutant males had significantly improved survival compared to control genotype (**Figure 3C**) but not in RNAi experiments (**Figure 3C’**). Therefore, an opposite survival trend in the null mutant flies would only be likely if the SNP was a gain-of-function mutation, which is difficult to test. Similarly, for the SNP (X_14160126) in gene *CG13404* (effect = 0.1213) predicts less susceptibility but we observed that RNAi knockdown females were significantly more susceptible compared to control genotype (**Figure 3E’**), while null mutants had no difference in host survival during infection (**Figure 3E**). Gene product threshold effects are one possible explanation for these complex data on host survival, and overall susceptible or tolerant phenotype during infection must be tested at the host level with subsequent functional experiments must be conducted.

Our screen using the DGRP identified several gene variants that affected host survival during *C. burnetii* infection that taken together with their known function and human ortholog information, can drive new mechanism-driven questions. Importantly, this study builds on our previously developed framework utilizing the *D. melanogaster* as an animal model to dissect the innate immune response to *C. burnetii* infection (Bastos et al., 2017; Hiroyasu et al., 2018). We observed that for candidate and validated genes, regulatory element data helps explain how the gene variants may affect host survival. In each case, identification of alternate transcripts in RNA-seq data would be supportive of the functional hypotheses suggested by our bioinformatic analyses. Follow up studies will need to go beyond whole organism mutants tested here, such as testing minigene mutants which allow researchers to test splicing patterns *in vitro* (Cooper, 2005; Ruan et al., 2015; Stoss et al., 1999). Nevertheless, the gene variants identified here highlight conserved cross-species connections and an opportunity for novel discovery about their role during *C. burnetii* infection.

## ACKNOWLEDGEMENTS

We thank Marcos A. Perez for critical review of this manuscript. We thank Michael D. Knight, Sarah A. Borgnes, Olivia Hayden, Marina Martin, and Emily L. Kindelberger for assistance in injecting fruit flies. We thank Codie Durfee for technical assistance. We are thankful to the Drosophila Genomics Resource Center (P40OD010949), the Bloomington Drosophila Stock Center (P40OD018537), the Vienna Drosophila Stock Center, Exelixis and TRiP at Harvard Medical School (R01GM084947) for providing reagents and fly stocks. This investigation was supported by funds from Washington State University and the National Institutes of Health Public Health Service grant R21AI128103 (to A.G.G.). R.D.O. was supported by NIH Training Grant T32AI007025. This investigation‘s contents are solely the responsibility of the authors and do not necessarily represent the official views of the NIH.

